# Safety Assessment of Broadband Laser Exposure in µOCT

**DOI:** 10.1101/2021.08.12.455289

**Authors:** Graham L. C. Spicer, Benjamin Child, Joseph A. Gardecki, Aditya Kumar, Andreas Wartak, Abigail Gregg, Hui Min Leung, Guillermo J. Tearney

## Abstract

Micro-optical coherence tomography (µOCT) improves the spatial resolution of *in vivo* OCT imaging by utilizing sophisticated focusing schemes and broadband illumination. This study explores the safety of coronary and trachea tissue exposure to µOCT illumination. © 2021 The Authors

## 1. Introduction

Micro-optical coherence tomography (µOCT) is a novel, high-speed optical imaging technology with high lateral and axial resolution on the order of microns [1]. For application to intravascular imaging, µOCT catheter technology has been developed which uses a novel focusing scheme to produce a coaxially focused multimode (CAFM) illumination profile to simultaneously extend the beam depth of focus (DOF) and narrow its minimum width along the DOF [2,3]. This differs from a traditional Gaussian beam used for catheter imaging with regard to the modal extent of optical power along the beam path and has the potential to locally concentrate power delivered to a sample due to constructive interference within the pseudo-Bessel beam profile [4]. The focused power distribution may therefore differ from that of the current clinical standard of care for intravascular and airway OCT catheters (NIR, spot size tens of micrometers) [5,6]. Hence, it is important to characterize any light-induced tissue damage that may occur with CAFM µOCT imaging at clinically relevant sample powers. While standards for maximum permissible exposure exist for skin and eye, they have not been reported for other tissues and may not be generally applicable for all illumination schemes [7]. Here we present the results of a study to validate the safety of the focusing schemes used in probe-based CAFM µOCT imaging for a range of broadband illumination powers and exposure durations.

## 2. Materials and Methods

The experimental setup for safety testing is shown in Fig. 1. Broadband (510-2400 nm) light from a supercontinuum laser (NKT Photonics SuperK Extreme EXR-15) was spectrally filtered to achieve a flattened spectrum from 650-950 nm with a dichroic mirror (DM) and custom shaping filter (SF), attenuated, and coupled into single mode fiber for delivery to the µOCT test probe. For all experiments, supercontinuum laser power was set to 100% and power was tuned by a partial shutter (PS) placed after free space filtering of the beam. Test probes were designed in Zemax OpticStudio to provide optimal focus at a working distance of 1.6 mm, constructed using established methods [2], and profiled in the focal region with a CMOS array and 9x relay lens (DataRay, Inc). Power out of the probe was measured with a thermal power meter (Thorlabs S415C, PM100D) with output normalized by the detector’s spectral response and the probe output spectrum, which was independently acquired with an optical spectrum analyzer (Anritsu MS9740B). The probe was mounted in a downward orientation to a motorized linear translation stage for the exposure protocol.

**Figure 1.**
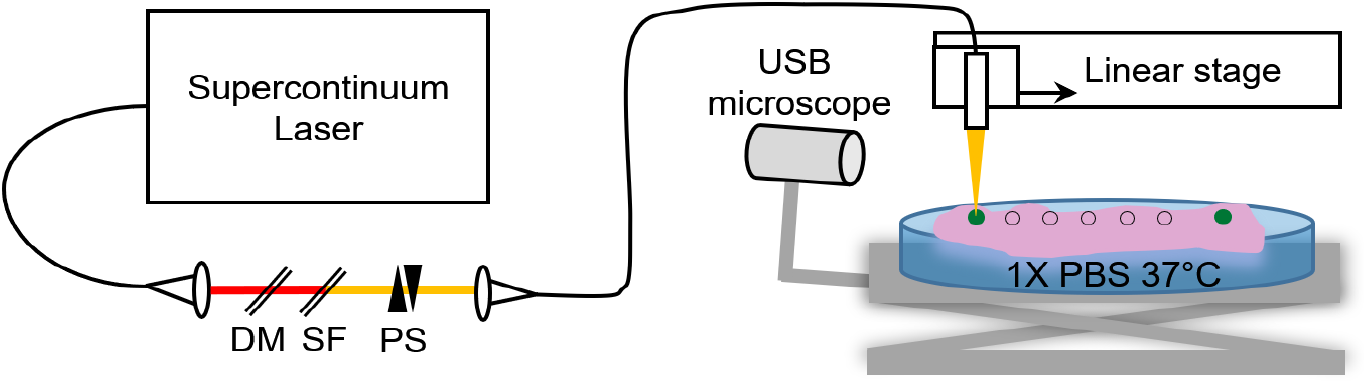
Schematic of experimental setup.

Fresh coronary arteries and tracheas were obtained from 75 kg swine (N=3) within one hour of animal sacrifice (IACUC protocol 2007N000041), after which the dissected tissue samples were kept in phosphate buffered saline (PBS) held at 37°C. Seven discrete exposures were generated along a linear path, with spacing 2 mm between the first six sites and 3 mm between sites 6 and 7. The proximity of the probe to the tissue surface was measured in real-time with a USB microscope and the tissue height was adjusted between each exposure site to place the probe’s focus at the tissue surface. Green tissue marking dye was applied to sites 1 and 7 prior to exposure to create a positive control burn site due to the increased optical absorption of the dyed tissue. After the exposure protocol was completed, the tissue was photographed, frozen at -20°C for 24 hours, then at -80°C for an additional 24 hours prior to embedding in optical cutting temperature medium for cryosectioning and histological processing. Tissue was sectioned through the visible green ink capturing 7 μm thick sections every 200 μm between levels. Frozen sections were then stained with nitro blue tetrazolium chloride (NBTC) for lactate dehydrogenase [8]. NTBC-stained slides were imaged with an automated slide scanner (Hamamatsu Nanozoomer) and inspected for laser-induced tissue damage, indicated by regions absent of blue stain.

## 3. Results

Truncated GRIN lens-based fiber probes with Gaussian and CAFM focusing schemes were designed to focus at a working distance of between 1 and 2 mm from the distal 0.5 mm diameter GRIN lens surface, a typical working distance and optic size used for intracoronary imaging. Probe designs and respective beam profiles are shown in Fig. 2.

**Figure 2.**
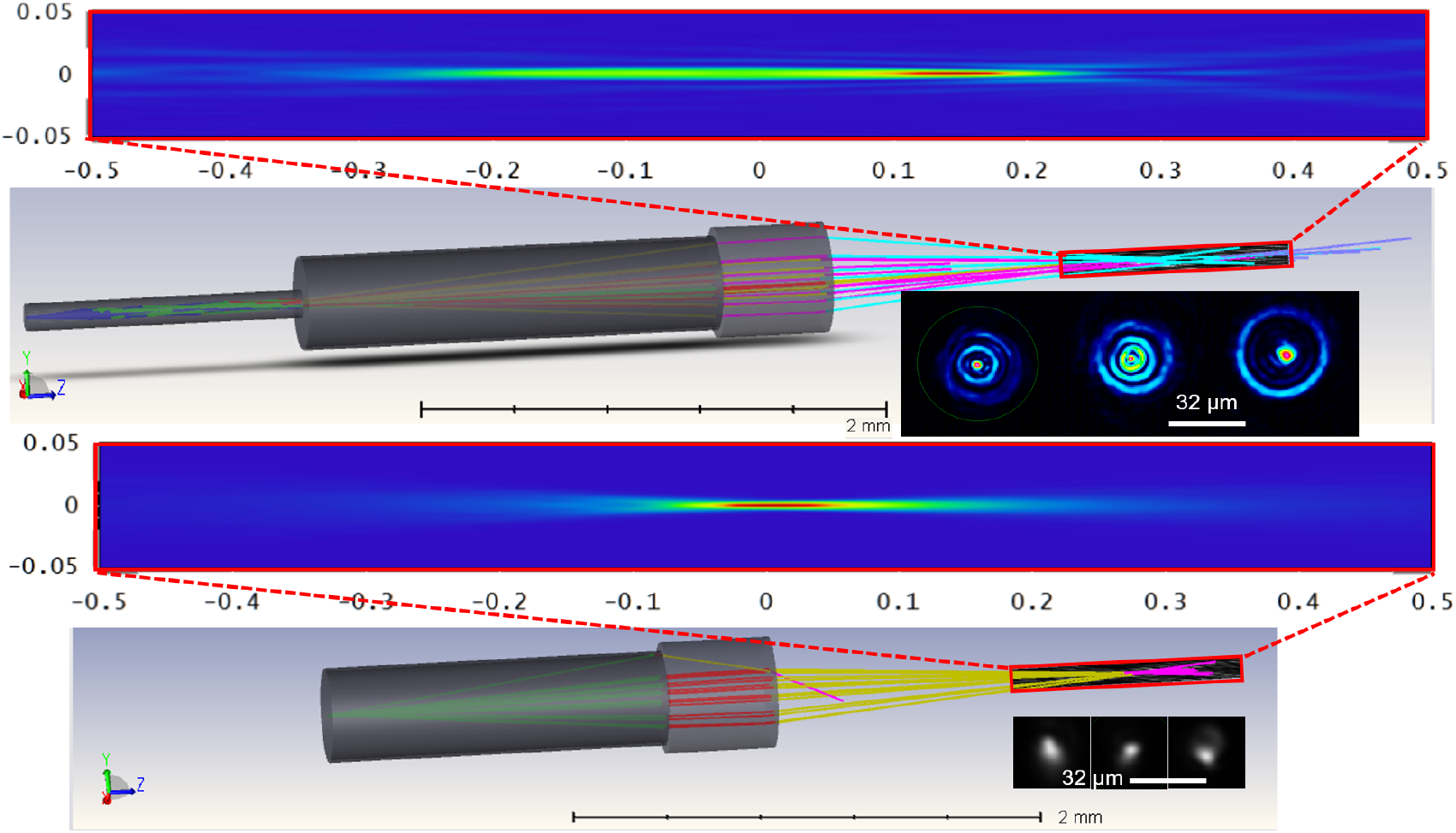
Design, simulation, and characterization of CAFM (top) and Gaussian (bottom) test probes, scale in mm.

In simulation and experiment, when compared with a Gaussian focus probe (no mirror tunnel), the CAFM probe produced tighter focus and an extended DOF, corroborating prior simulation and experimental investigation [3]. The potential for laser-induced damage was evaluated for both OCT illumination schemes in duplicate samples (n=2) of coronary artery and tracheal tissue for a range of exposure times and powers summarized in Table 1.

**Table 1.**
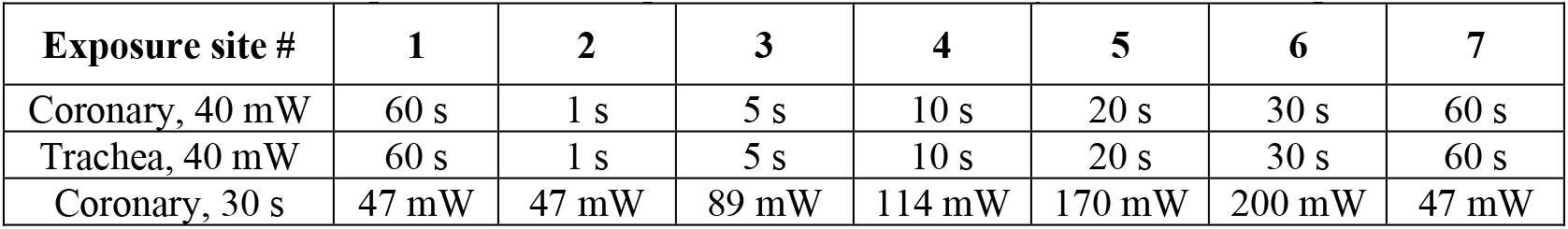
Exposure times and powers tested on coronary and trachea samples.

| Exposure site # | 1 | 2 | 3 | 4 | 5 | 6 | 7 |
| --- | --- | --- | --- | --- | --- | --- | --- |
| Coronary, 40 mW | 60 s | 1 s | 5 s | 10 s | 20 s | 30 s | 60 s |
| Trachea, 40 mW | 60 s | 1 s | 5 s | 10 s | 20 s | 30 s | 60 s |
| Coronary, 30 s | 47 mW | 47 mW | 89 mW | 114 mW | 170 mW | 200 mW | 47 mW |

Tissues were sectioned at 200 μm intervals which provided adequate sampling to detect positive control burn damage at exposure sites 1 and 7 across multiple slides, with damage regions extending laterally >300 μm for all slides where positive control burns were apparent. An example photo of a coronary sample prior to freezing with exposure sites and region where histology was taken is shown in Fig. 3A. Representative NBTC histology images of an enlarged positive control burn site, and coronary and trachea slices through the exposure plane are shown in Fig. 3B-D, respectively.

**Figure 3.**
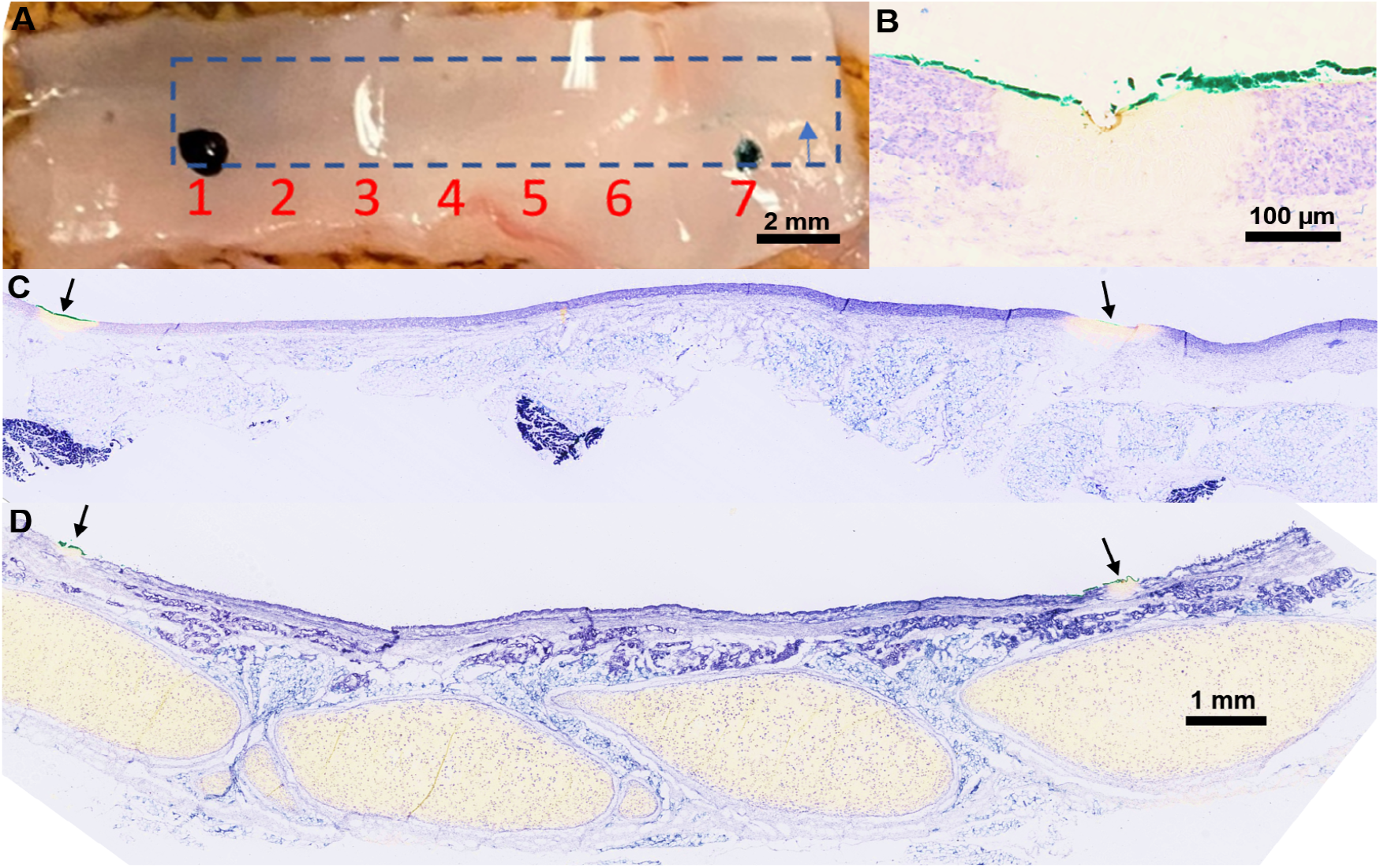
Representative photo and histology of test samples. Blue box in (A) indicates section of tissue processed for histology, with blue arrow indicated direction of subsequent slicing planes into tissue block. Black arrows in (C) and (D) indicate exposure sites 1 and 7 (positive control).

For all test samples, laser-induced damage was observed at the positive control exposure sites 1 and 7, but no damage was observed for any other exposure site, suggesting that the combination of exposure parameters and illumination scheme (pattern) will maintain tissue viability after *in vivo* scanning.

## 4. Discussion

In this study we demonstrate the general safety of Gaussian beam and CAFM beam focusing schemes for the broadband (650-950 nm) illumination used in µOCT probes for high resolution intravascular and pulmonary imaging applications. CAFM beam optics previously described for OCT lateral resolution and depth of focus enhancement, while providing tighter focus for select axial positions along the beam, do not induce tissue damage in this study. While simulation indicates the power density at the central axis of the CAFM beam may reach levels on the order of 3×10^4^ mW/cm^2^ for a 40 mW total power input, the power is also divided among multiple focal modes which lessen the total energy delivery at the tight focal regions, effectively smearing energy delivery along depth. While this study demonstrates the safety of static illumination of vascular and pulmonary tissues with OCT focusing schemes, imaging is achieved through rapid scanning of the beam across tissue samples which reduces dwell time within a single focal volume to the order of microseconds. In conclusion, we anticipate that higher exposure safety limits for µOCT imaging suggested by this study will permit development of faster, micron-level resolution intravascular imaging catheters that achieve adequate sampling for clinical use.

## 5. Acknowledgements

The authors acknowledge funding for this work from Verdure Biotech Inc. and histology services from the Wellman Center for Photomedicine Photopathology Core. This work was presented at European Conferences on Biomedical Optics OSA/SPIE (ECBO 2021), June 22^nd^ 2021, Munich, Germany. Paper ETu4A.3

